# Proper modelling of ligand binding requires an ensemble of bound and unbound states

**DOI:** 10.1101/078147

**Authors:** Nicholas M Pearce, Frank von Delft

**Affiliations:** Structural Genomics Consortium, University of Oxford, Roosevelt Drive, Oxford, OX3 7DQ, UK.; Diamond Light Source Ltd, Harwell Science and Innovation Campus, Didcot OX11 0QX, UK; Department of Biochemistry, University of Johannesburg, Auckland Park, 2006, South Africa

## Abstract

**Synopsis:** We emphasise and demonstrate the importance of modelling the superpositions of ligand-bound and unbound states that commonly occur in crystallographic datasets. Generation of an ensemble that describes not only the dominant state in the crystal is important for the high-quality refinement of low-occupancy ligands, as well as to present a model that explains all of the observed density.

**Abstract:** Small molecules bind to only a fraction of the proteins in the crystal lattice, but occupancy refinement of ligands is often avoided by convention; occupancies are set to unity, assuming that the error will be adequately modelled by the B-factors, and weak ligand density is generally ignored or attributed to disorder. Where occupancy refinement *is* performed, the superposed atomic state is rarely modelled. We show here that these modelling approaches lead to a degradation of the quality of the ligand model, and potentially affect the interpretation of the interactions between the bound ligand and the protein. Instead, superior accuracy is achieved by modelling the ligand as partially occupied and superposed on a ligand-free “ground-state” solvent model. Explicit modelling of the superposed unbound fraction of the crystal using a reference dataset allows constrained refinement of the occupancy of the ligand with minimal fear of over-fitting. Better representation of the crystal also leads to more meaningful refined atomic parameters such as the B-factor, allowing more insight into dynamics in the crystal. We present a simple approach and simple guidelines for generating the ensemble of bound and unbound states, assuming that datasets representing the unbound states (the ground state) are available. Judged by various electron density metrics, ensemble models are consistently better than corresponding single-state models. Furthermore, local modelling of the superposed ground state is found to be generally more important for the quality of the ligand model than convergence of the overall phases.

## 1. Introduction

Crystallographic diffraction experiments are used to reveal the atomic composition of protein crystals, but where the crystal is composed of objects in multiple states, the resulting diffraction pattern is a weighted average of these states. Ligands will often – and likely almost invariably – bind at sub-unitary occupancy; the subsequently derived electron density consists of an average over the bound state and the corresponding unbound state (which we term the *ground state*). However, it is standard practice not to model a superposition of multiple states, but instead to model only the ligand-bound conformation (and furthermore normally at unitary occupancy); this is commonly observed in the PDB^1,2^ (Figure 1).

**Figure 1.**
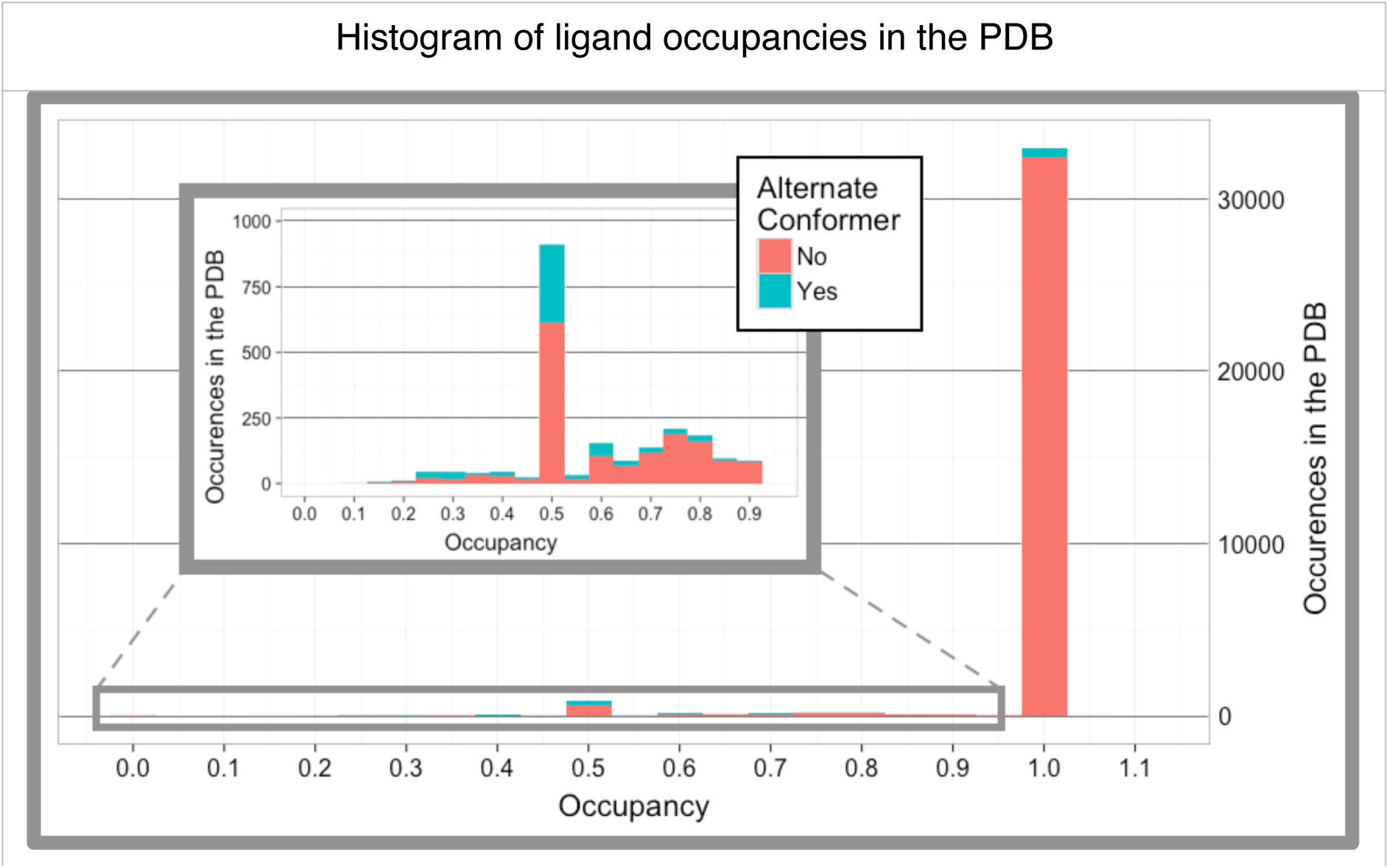
Most ligands in the PDB are modelled at unitary occupancy, and many partial occupancy ligands are not modelled with an alternate state. Histogram of all ligand occupancies in the PDB classified by the presence of an alternate conformer identifier (red: no conformer ID, blue: modelled with a conformer ID). Sub-unitary occupancy ligands are clarified in the inset graph. Only the first instance of each ligand type from each PDB structure was used; following this all ligands with fewer than 5 non-hydrogen atoms and more than 50 instances were removed to avoid bias towards common molecules. Where alternate conformations of ligands are present, the total occupancy is used. The large majority of ligands are modelled at unitary occupancy (32396, 92.1%). A smaller number have non-unitary occupancies but no alternate conformer identifier (1640, 4.7%). The remainder are modelled using alternate conformers (1122, 3.2%), of which 548 are ligands with alternate conformers that sum to unitary occupancy. Worryingly, there are also ten instances with more than 100% occupancy. These modelling statistics are unlikely to represent the true situation in crystal structures, where ligands will rarely bind at near-full occupancy; ligands will always have a superposed solvent model where present at partial occupancy.

Occupancy refinement of ligands is likely avoided due to well-known interdependencies, instabilities and ambiguities that can occur in the simultaneous refinement of both B-factors and occupancies: improvements in crystallographic model fit can equally well be achieved by reducing occupancy or increasing B-factors^3^. When ligands are modelled at full occupancy, any resulting error is absorbed by inflating the refined B-factors. One is led to conclude that occupancy refinement is only deemed necessary when difference density appears over the ligand model, an impression corroborated by multiple conversations in online discussion fora such as *ccp4bb* and *ResearchGate*.

If occupancy refinement of the ligand-bound state is performed without a superposed solvent model, this implicitly implies that the rest of the crystal is either represented by vacuum – which is highly unlikely – or by bulk solvent, depending on the refinement program used. Close to the surface of the protein, it is unlikely that the solvent is truly represented by a bulk solvent model; this is especially true of binding sites, where solvent and buffer molecules will often bind in an ordered fashion at high occupancy, as in the examples presented in section 4. The absence of a superposed solvent model is a glaring modelling omission, and here we set out to show that inclusion of the superposed unbound state not only leads to a more complete model of crystal, but to a higher quality ligand model.

## 2. More complete models through explicit inclusion of the ground-state

We propose that ligands will – in the general case – *always* be better modelled with explicit representation of the superposed solvent state, determined from a ground-state crystal of the protein. Inclusion of the ground-state allows the occupancy of the superposed states to be constrained in refinement, reducing the ambiguity from simultaneous refinement of b-factors and occupancies.

This approach requires a credible model of the ground-state to be available. This is indeed the case in a large proportion of ligand-binding experiments, where ground-state (ligand-free) crystals are easily obtained, e.g. experiments where ligands are “soaked” into pre-formed crystals. Where ground-state crystals are difficult to generate, *e.g.* where the ligand stabilises a particular protein conformation and thus crystal form, the assumption of an ensemble is in any case unlikely to be relevant.

Once the ground-state structure of the protein has been determined, the corresponding atoms can be directly transferred to the model of any subsequent dataset of the same crystal form. Specifically, the ground-state model is combined with the changed-state (ligand-bound) conformation, and refined as an ensemble. Generating this ensemble is algorithmically simple for datasets that are reasonably isomorphous; where this is not the case, the unbound structure would require local alignment of corresponding atoms, although methods to do this robustly do not currently exist, to our knowledge.

In-between cycles of reciprocal-space refinement – if the crystal system is highly isomorphous – the ensemble model can be modelled or visually validated in programs such as Coot^4^ by alternating between real-space refinement of the ground-state model into a ground-state map (left-hand column, Figure 2), and checking the validity of the complete model in the ligand-bound dataset. In the case of a PanDDA-determined model^5^, additional maps are available for the modelling of the ligand-bound conformation (right-hand column, Figure 2). The PanDDA implementation further performs automatic merging of the changed-state model and the ground-state model, allowing ensembles to be utilised with little additional effort.

**Figure 2.**
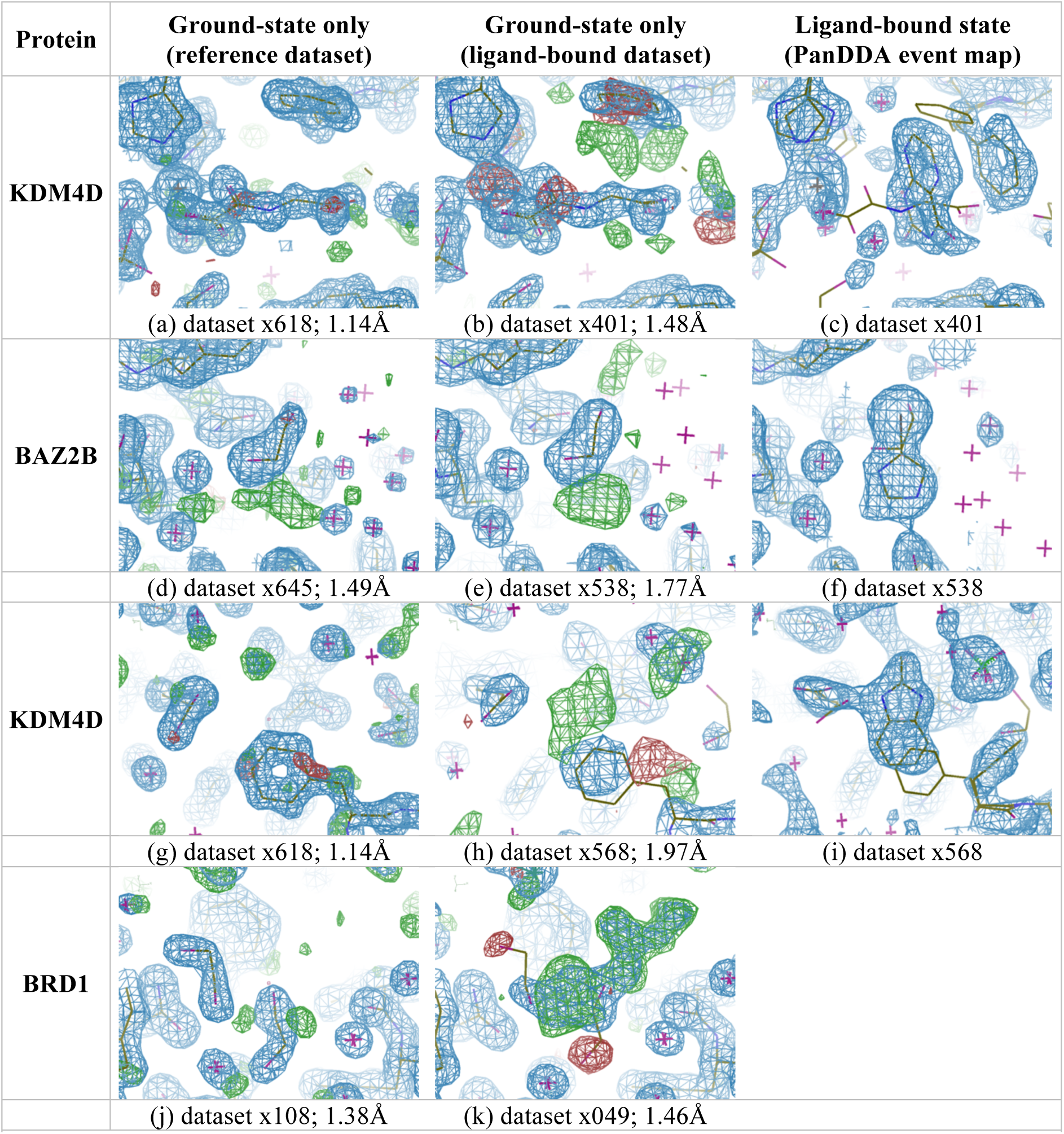
Determining the different crystal states requires different datasets. First two columns: 2mF_O_-DF_C_ maps contoured at 1.5σ (blue) and mF_O_-DF_C_ maps contoured at ±3σ (green/red). Last column: PanDDA event maps (blue) contoured at (c,f) 2σ or (i) 1σ. Resolutions are as indicated. <u>First column:</u> A reference dataset provides the ground-state model of the crystal. <u>Centre column:</u> The groundstate refined into a ligand-bound dataset leaves (generally uninterpretable) residual density for a superposed state. <u>Last column:</u> The PanDDA event map provides clear density for the ligand-bound model of the crystal (the superposed ground-state model is shown for reference). (a-c) Example from section 4.1. (d-f) Example from section 4.2. (g-i) Example from section 4.3. (j-k) Example from section 4.4; the event map is not shown since it is not required.

During modelling and refinement, the ground-state model should be considered a Bayesian prior, such that the underlying ground-state structure is assumed not to change from crystal to crystal. This applies even if the ground-state is not clearly discernible in the electron density; minor states will be “masked” by superposed major states, but they will still remain except where the ligand is truly unitary occupancy. In cases where the ground-state structure is crystallographically ill-defined in the ligand-bound-dataset (such as at low-resolution) it may be necessary to restrain the ground-state model to the reference dataset during refinement^6–8^.

This restraint addresses the main risk inherent in ensemble approaches, namely over-modelling the observed density by including additional, unwarranted atoms: including the ground-state model has a strong, first-principles rationale, and the information is derived from independent measurements. While interpreting the remaining density may not be easy in general, methods such as PanDDA^5^ address this problem explicitly by deconvoluting the superposition.

### 2.1. Systematic labelling of multiple crystal states to maximise interpretability

Locally heterogeneous crystal states are modelled through the use of alternate conformers, which ascribe each atom to a particular state of the crystal. Only for completely independent ensembles of models are alternate model identifiers utilised^9^. When merging the ground-state of the crystal with the ligand-bound state, the same conformer ID – sometimes referred to as the *altloc* or *altid* – should be given to all atoms of the same state. Each state may then be extracted by selection of a particular conformer from the ensemble, enabling the use of the structure by non-crystallographers; the superposed ground-state is essentially an experimental artefact. The occupancies of the different states may further be grouped during refinement, and the occupancies of the states constrained to sum to unity.

The clearest interpretation of the model is achieved when conformers are used for the bound and unbound states that are not used elsewhere in the structure; this prevents potential association of similarly-labelled alternate conformers that are causally unrelated. In the case of a single conformer for each bound/unbound state (where alternate conformers elsewhere in the structure, unrelated to binding, are A and B only), all ground-state-only atoms may be set to conformer C, and all bound-state-only atoms may be set to conformer D. This assignment of logical conformer IDs is automatically performed during the merging of the ground-state and the ligand-bound state within the PanDDA implementation; this automation greatly simplifies the modelling process, where the ground-state model is used as the starting model for analysis.

However, the limitations of alternate conformers can quickly manifest themselves where multiple conformations are present in the bound/unbound states. Since alternate conformers do not support branching of conformations (where e.g. an alternate conformation of the backbone can have two sidechain conformers), it may be necessary to introduce redundant alternate conformations for single-conformer residues to create contiguous models (Figure S1).

### 2.2. Local model completeness versus overall phase quality

Conventional crystallographic dogma states that high quality (near-convergence) phases are needed for the “optimal” crystallographic model to be obtained. However, we show in this work that the current convention of omitting the superposed unbound state is more detrimental to the quality of the ligand model than the degradation of the overall model phases. To compare the effects of global phase degradation, a “degraded-phase” model is produced in each of the examples in section 4. We begin with the final “optimal” model – where the ligand is modelled in superposition with a ground-state model – and distort the structure of the protein in regions distant from the ligand binding site, thereby introducing global phase error. Induced mean model phase difference relative to the full ensemble model is in the range of 20-30° (as calculated by cphasematch^10^). Further details may be found in section S1.

## 3. Qualitative and quantitative comparison of different modelling approaches

To demonstrate the improvement of ligand models through inclusion of the superposed ground state, we present four examples in section 4, covering a range of ligand occupancies. All ligands were identified with PanDDA Z-maps and ligand-bound states were modelled using the PanDDA event maps. Three models of the crystal containing ligands are refined and compared: a ligand-state-only model; a high-quality ensemble model; and a degraded-phase ensemble model. A solvent-state-only model is also refined for completeness (central column, Figure 2).

The ligand-state-only model for refinement is obtained by removing the ground state from the ensemble and setting the ligand occupancy to 0.95. The solvent-state-only model is similarly generated by removing the ligand-bound state and setting the solvent occupancy to 1.0 (this simulates the normal modelling case, where the solvent occupancy would not typically be refined). Degraded-phase models are created from the ensemble models as described in section 2.2. All models are refined with phenix.refine^11^ (version 1.9-1682) using the default parameters against crystallographic data from before a ligand was placed, to prevent phase bias. Ligand occupancy is refined for all models; for the ensemble models, the occupancies of superposed states are constrained to sum to unity.

### 3.1. Utilisation of validation metrics for quantitative model comparison

The refined ligand models are compared using a variety of density- and model-based validation metrics; these metrics and their optimal values are described in Table 1. Density metrics – all calculated by EDSTATS^12^ – include the conventional real-space correlation coefficient (RSCC), but also newer metrics such as RSZD and RSZO^12^. Tickle (2012) shows that these new metrics can be used to ask more detailed questions about the model: RSZD measures the accuracy of the model through the analysis of difference density, highlighting modelling errors, and RSZO measures the precision of the density for the model, highlighting weak features. RSZO is calculated by taking the average of the density over the model and dividing by the noise in the map; since the amount of density for a residue is directly related to the occupancy of the residue, we divide RSZO by the occupancy of the residue to give a normalised value (RSZO/OCC) that can be used to compare models at different occupancies in the same dataset. We also calculate the B-factor ratio of the ligand to the surrounding protein residues (within 4Å) to measure the consistency of the ligand model with its local environment; as well as the RMSD of the refined ligand and the fitted ligand, to measure the (in)stability of the model coordinates in refinement. These measures are displayed visually as radar plots, where the “better” the metric value, the closer it is to the centre of the plot. The axes of the radar plot are scaled such that the “best” value is plotted at the centre of the plot and the “worst” value is plotted at the extreme of the axis.

**Table 1.** Electron density and model metrics used for the validation of crystallographic models. The combination of five metrics highlights a variety of features of models, and together allow for a comprehensive description of the atomic model of a residue. RSCC ensures good overall similarity of the model to the density. RSZD measures the difference density over the model, highlighting errors or the presence of currently un-modelled or over-modelled atoms. RSZO indicates density strength, and the normalisation by occupancy can indicate errors in the occupancy of a model or a misplaced or absent model. The B-factor ratio highlights errors in the B-factors of a residue, as these should be consistent with its surroundings: physically, there cannot be step changes in mobility of atoms in a crystal. The RMSD measures the movement of residues in refinement; a numerically unstable residue may be indicative of error in the model. All density metrics are calculated using EDSTATS^12^.

| Metric | Description | Preferred values |
| --- | --- | --- |
| RSCC | Correlation between model and observed electron density | > 0.7 |
| RSZD | Statistical measure of difference density in region of model | < 3 |
| RSZO/OCC | Strength of density over model, normalised for occupancy | > 2 |
| B-factor Ratio | B-factor ratio of residue atoms and sidechain atoms within 4Å | ~ 1 |
| RMSD | Root-mean-squared-deviation of the atomic coordinates | < 1 |

#### 3.1.1. Effects of phase quality on model validation metrics

The RSZD metric is less informative when analysing models with poor phases, because it is dependent on the quality of the model phases. RSZD and RSZO are derived with the assumption of near-convergence phases, and use an estimation of the noise in the maps to calculate quality criteria for residues. Lower RSZD would normally indicate a better model, but this is not the case here: when the quality of the phases is reduced, the noise in the maps also increases, and therefore decreases both RSZD and RSZO, regardless of whether the model has changed.

## 4. Results

We now present several cases where the inclusion of a complementary solvent model leads to a better description of the crystal, and thereby a higher-quality ligand model. The models here were all identified and modelled using the PanDDA method^5^. The model of the ligand was in each case derived from PanDDA “event” maps, and we investigate here only the effect that the inclusion/absence of the superposed solvent model has on the interpretation of the data. Models are generated and refined as described in previous sections. Validation metrics are calculated for only the ligand residue in each of the models. Crystallographic model parameters, including ligand validation scores, may be found in section S1. Details for obtaining the crystallographic data can be found in the PanDDA publication^5^.

### 4.1. Binding of the ligand across a bound substrate mimetic

To demonstrate the process of modelling both states, we first present an example where a strongly bound substrate mimetic is superposed with a weakly-bound soaked ligand, and an ensemble is clearly necessary. N-oxalylglycine (NOG) is tightly bound at high occupancy (~90%) in the ground-state crystal form of human Lysine-specific demethylase 4D (KDM4D), as shown in the reference dataset (Figure 2a). A soaked ligand binds across this substrate mimetic in a small fraction of the crystal, as shown in the PanDDA event map (Figure 2c). Modelling of the two states can thus be performed separately, and merged for refinement; when refined as an ensemble, the superposition of the two states leads to a good model, with negligible amounts of difference density remaining (Figure 3b).

**Figure 3.**
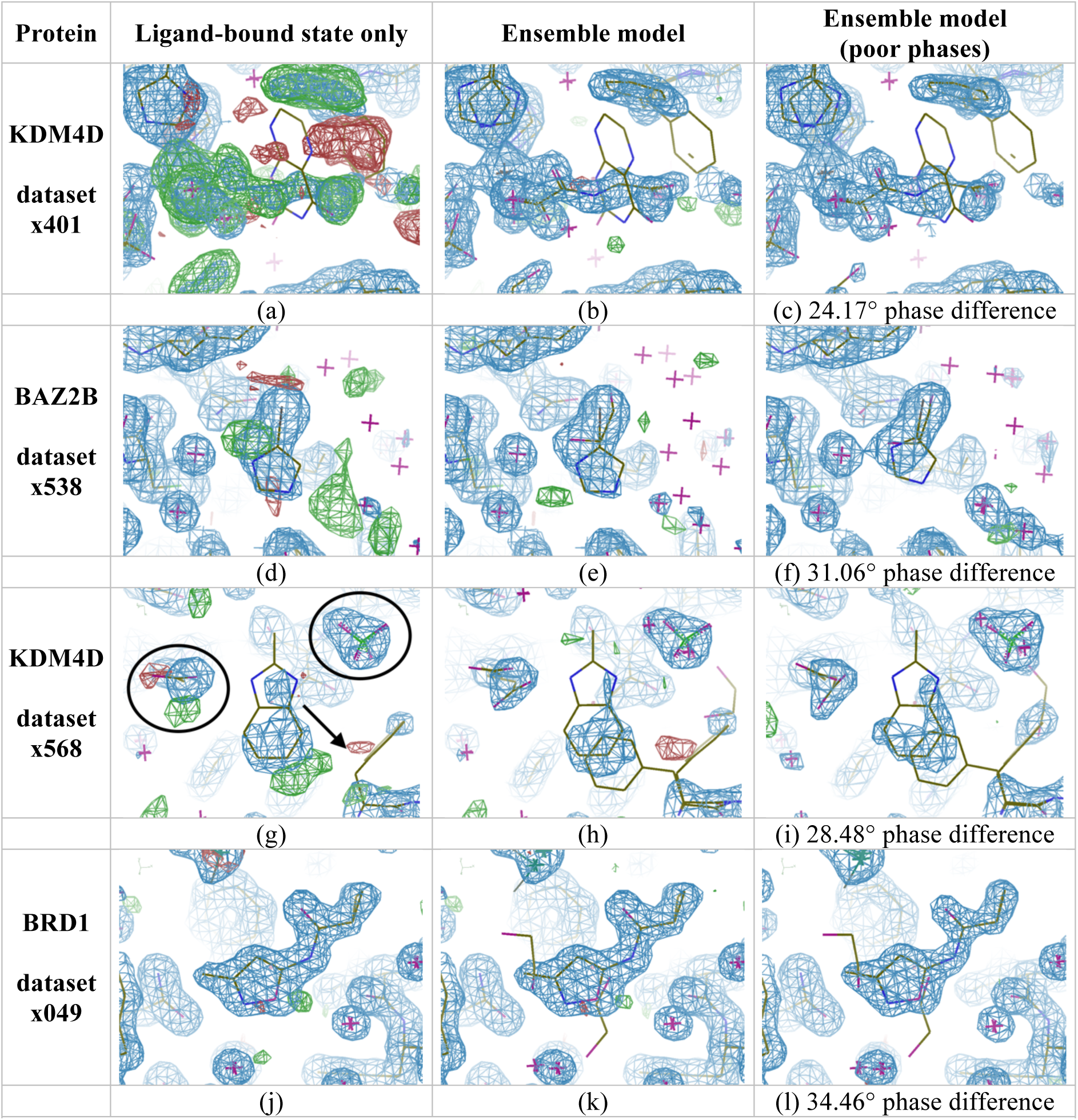
Ensemble models consistently leave less residual difference density than ligand-only models. All images: 2mF_O_-DF_C_ maps contoured at 1.5σ (blue) and mF_O_-DF_C_ maps contoured at ±3σ (green/red). <u>First column:</u> Refinement with the ligand model only. <u>Centre column:</u> Refinement of the crystal as an ensemble of states. <u>Last column:</u> Refinement of the crystal as an ensemble of states with a degraded protein model (phase difference as indicated, relative to the ensemble model). (a,d,g) Modelling the ligand but removing the ground-state leads to difference density for the absent state, and in (d) the ligand moves into density for the ground-state. (j) Removing the ground-state for a high occupancy ligand (refined value 0.89) does not lead to discernible difference density. (b,e,h,k) Refinement of ensemble models explain all of the observed density, and ligands do not move from the fitted pose (confirmed by the validation plots in Figure 4). (c,f,i,l) Refining with degraded phases leads to only minor visual differences, except in (f) where the ligand moves relative to the fitted pose.

Although not interpretable, residual difference density can still be seen for the bound ligand when the ground-state model is refined alone (Figure 2b). As expected, refinement of the ligand without the superposed NOG results in a poor quality model (Figure 3a), because a large fraction of the crystal is locally unrepresented; refinement of the ensemble results in a better model for the ligand (Figure 3b), scoring well across all 5 metrics. On the radar validation plot (Figure 4a) this is shown as the ensemble-model line (green) being entirely contained within the ligand-only line (red) – the closer the line is to the centre of the plot, the better the model. Optimal modelling of the ligand requires the superposed ground-state conformation to be present in refinement.

**Figure 4.**
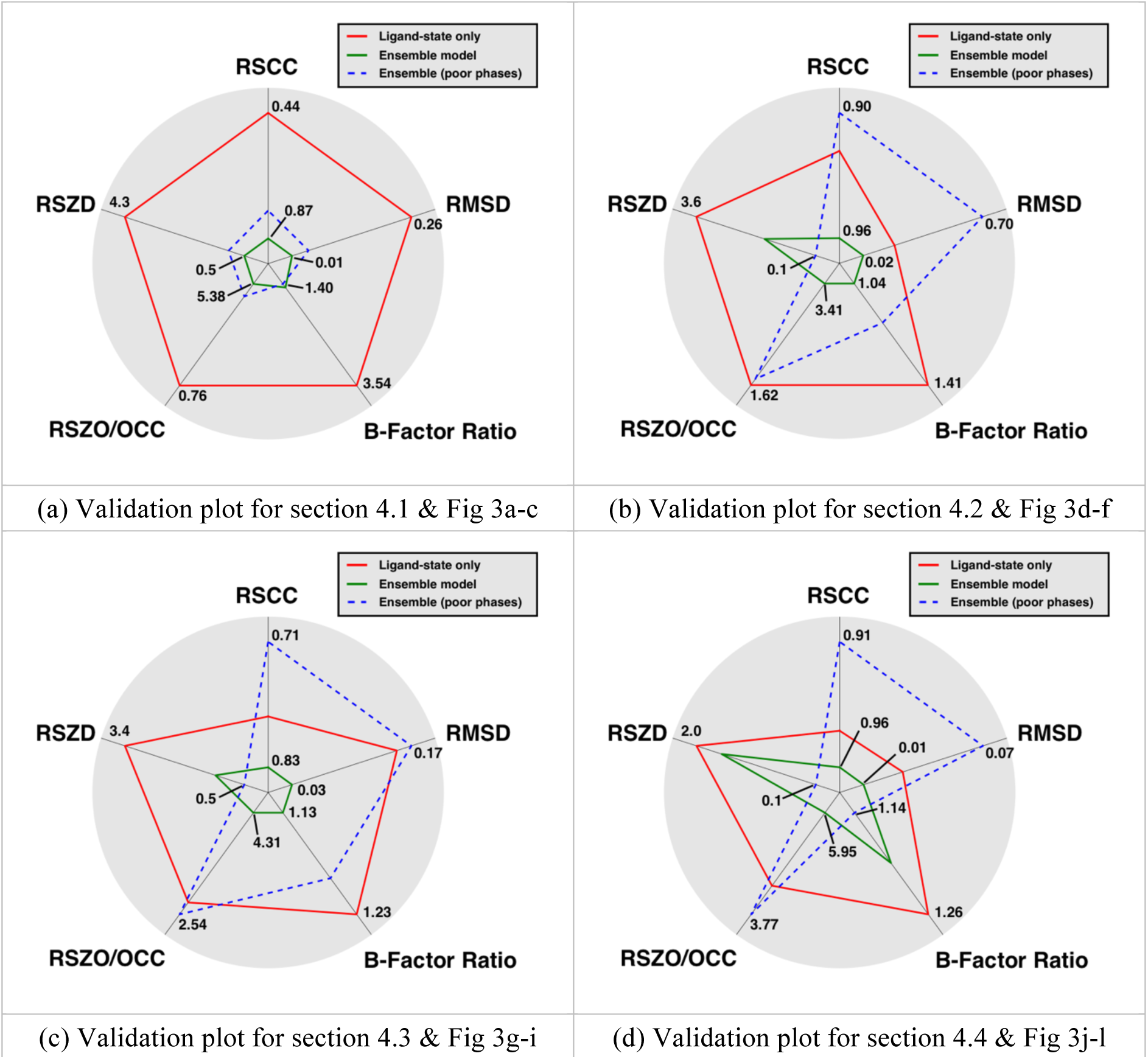
Validation plots for the different modelling approaches: axes are not absolute, but have been scaled relative to minima and maxima of the plotted values, and only the minimum and maximum values are marked on the axes; for all model scores refer to section S1. (a) Plots for Figure 3a-c. The plot confirms the visual inspection of the electron density; the ligand scores are improved across all metrics when refined as an ensemble, relative to the ligand modelled alone. The absence of the superposed substrate model has a greater effect on the ligand model than the degradation of the protein model phases. (b) Plots for Figure 3d-f. The ensemble model provides the best model for the ligand. The RSZD is decreased in the degraded-phase model for reasons explained in the main text, and is not related to an improved model. (c) Plots for Figure 3g-i. Once more, the model statistics are improved with the addition of a superposed solvent model, with the caveat that the lower RSZD for degraded phases is not indicative of an improved model. (d) Plots for Figure 3j-l. The inclusion of the solvent model still increases the quality of the model compared to when it is omitted, albeit marginally. The degraded phase model has lower B-factor ratios than either of the other two models due to a decrease in the B-factors of the ligand and a corresponding drop in occupancy.

The degraded protein model (Figure 3c) has a 31° average phase difference to the high-quality ensemble model, increasing the R-free from 17% to 29%. However, the model of the ligand is not significantly degraded, and still scores well on all five model validation metrics, although worse than the ensemble model with high-quality phases. In this case, the correctness of the local model is more important than the convergence of the global phases.

### 4.2. Binding of a ligand in place of a solvent molecule

In a soaked crystal of human Bromodomain Adjacent to Zinc finger domain 2B (BAZ2B), an ethylene glycol is bound in a semi-ordered fashion, with a superposed ligand, to the asparagine in the binding site. The solvent model derived from a reference dataset is not optimal, and some difference density remains even when a ligand is not present (Figure 2d). Refinement with the ground-state model in the ligand-bound dataset does not lead to significant additional difference density, as the refined solvent model masks the presence of the ligand’s bromine (Figure 2e).

The PanDDA map, however, shows clear evidence for the ligand (Figure 2f); the positioning of the bromine can also be confirmed by an anomalous difference map (not shown). Refinement with only the bound state causes the ligand atoms to be pulled into the density for the ethylene glycol, and difference density remains (Figure 3d). Refinement of the ensemble leads to a good model (Figure 3e), with all density well-explained, and no movement of the ligand from the fitted pose.

Refinement of the degraded-phase model (Figure 3f) also causes the ligand to move relative to the fitted position. In this case, the absence of the superposed model and the quality of the model phases are both important for the quality of the final ligand model, reflected by the validation metrics (Figure 4b).

It is noteworthy that the RSCC of the ligand in all models is greater than 0.9, showing that whilst a large RSCC is necessary for a good model, it is not sufficient to determine the quality of the model: it does not account for the presence of difference density. As explained in Section 3.1.1, the RSZD of 0.1 for the degraded-phase ligand model, which would normally indicate a very good model, is affected by noise in the maps from the degraded phases; the RSZD is very sensitive to the overall correctness of the model. Multiple validation metrics, as well as a near-complete model, are needed to validate weak features.

### 4.3. A binding ligand overlaps with alternate conformations of a sidechain

Another ligand in a KDM4D dataset binds along with a sulphate to a putative allosteric site. Refinement with the ground-state conformation leaves residual unmodelled difference density (Figure 2g,h). The pose and identity of the ligand is clearly revealed in the PanDDA event map (Figure 2i), revealing the re-ordering of two sidechains and that the ligand is superposed on the ground-state conformation of the phenylalanine.

Upon inspection of the refined ensemble model (Figure 3h), it was suggested to the authors by another experienced crystallographer that the ground-state conformation should be deleted and the ligand-bound state refined as the sole conformation. This recommendation supports our observation that the pervading convention – to generate only a single conformation of the crystal wherever possible – dominates even in the face of clear evidence that multiple states are present. The density in the area of overlap between the ligand and the phenylalanine is significantly stronger than over the rest of either residue, and difference density is present when either state is refined separately (Figure 2h, Figure 3g). The residual density from the ligand-state-only model (Figure 3g) might further tempt a crystallographer to move the model down and right by ~1Å (as indicated by the arrow in Figure 3g), although this causes clashes with the C_β_ of the phenylalanine and adversely affect the interactions that the ligand makes with the aspartate and the sulphate (marked with ovals in Figure 3g). All evidence points towards the presence of multiple states in the data, and therefore these multiple states should be present in the model.

The phase degradation in Figure 3i (mean phase difference to ensemble model 28.48°) degrades the ligand model RSZO and the B-factor ratio to a similar level as the omission of the ground state model, and significantly degrades the RSCC (Figure 4c). Again, we observe a decrease in RSZD with the decrease in phase quality. The ensemble model provides the best interpretation of the experimental data.

### 4.4. Traces of the ground state remain, even for a high occupancy ligand

One ligand screened against the bromodomain of BRD1 binds strongly in the principal binding site (Figure 2j,k), with a refined occupancy of 84-89% (multi-state and ligand-only refined occupancies respectively). In the reverse case of section 4.1, the ligand occupancy is much higher than the ground-state occupancy, and this ligand would conventionally be modelled at unitary occupancy.

Once more, inclusion of the ground-state solvent improves the model quality, although in this case only marginally (Figure 3j,k & Figure 4d). Even with this strong binder, visual traces of the ground-state model remain: contouring the 2mF_o_-DF_c_ map to zero rmsd shows some evidence for ground-state solvent (Figure 5).

**Figure 5.**
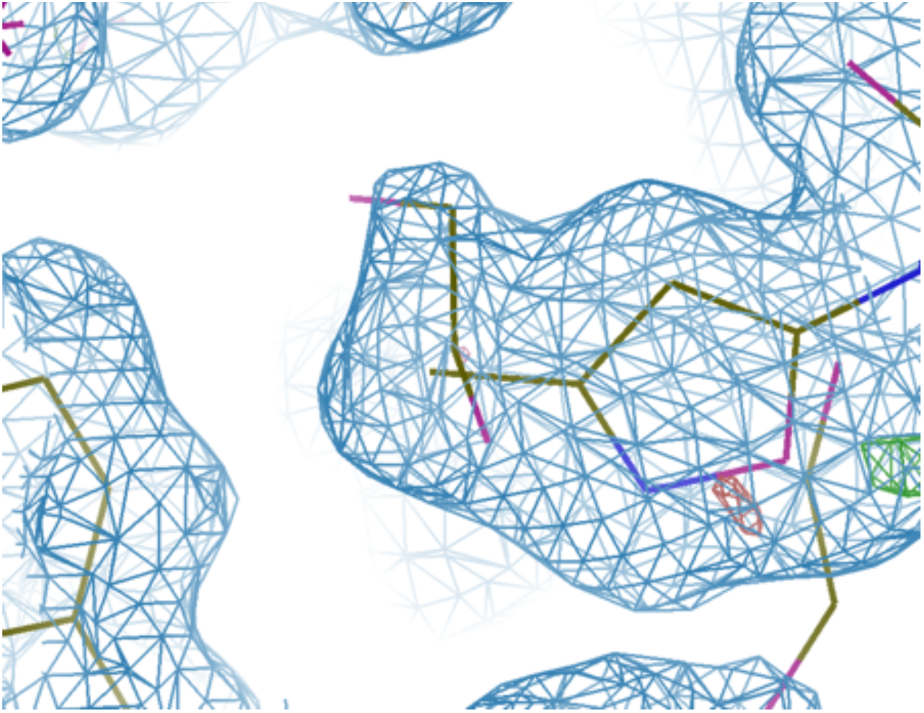
Weak density for the ground-state model is still visible in refined maps. No evidence is seen that would support the removal of the superposed solvent model. 2mF_o_-DF_c_ map (blue) contoured at 0.0σ, mF_o_-DF_c_ map (green/red) contoured at ±3σ.

Phase degradation degrades the RSCC, RMSD and the RSZO more than the absence of the solvent model, with a decrease in RSZD as previously. Here the B-factor ratio is seen to be lower for the phase-degraded model than for the other models, due to a decrease in the B-factors of the ligand by two, and a corresponding decrease in the occupancy to 0.77; this behaviour demonstrates the ambiguity that can be observed in simultaneous refinement of B-factors and occupancies.

## 5. Discussion

The examples presented here show that there is consistent evidence that ground-state molecules are superposed in the experimental data on top of binding ligands across a range of non-unitary occupancies. We have also shown that the inclusion of a superposed ground-state model, obtained from a reference dataset, improves the quality of obtained ligand models in all cases. In the case of some weak ligands, the ground state model is crucial for the refinement of the protein/ligand complex (section 4.1); in other cases it acts simply to remove “extraneous” difference density that could be interpreted by an over-zealous modeller as being caused by a ligand in multiple conformations (section 4.2). The modelling approach can affect the interpretation of inter-molecular interactions (section 4.3), and in the case of high occupancy, a superposed ground state can still marginally improve the ligand model, alongside providing a complete model of the crystal (section 4.4).

With the current increase in popularity of experiments such as fragment screening by crystallography amongst academic groups, the PDB is set to see a sharp increase in structures that contain binders with considerably less than unitary occupancy (e.g. [^13^]). We have shown that the models of such partial-occupancy ligands benefit from the inclusion of a superposed ground-state; from these results, we propose that a new standard modelling convention is adopted, where bound ligands are modelled as a superposition of states *wherever possible*. Experimentally this is no extra burden, as an unbound reference dataset is normally already available when soaking experiments are performed. Computationally, however, this will require the implementation of tools for the trivial generation of ensembles from multiple single-state models; the PanDDA implementation goes some way towards achieving this new paradigm.

Performed correctly, the addition of a solvent model allows no further degrees of freedom for the crystallographer, as the ground-state model is solely determined in an orthogonal reference dataset. Utilisation of prior knowledge in the modelling process will lead to higher quality crystallographic phases, and should ultimately contribute to closing the R-factor gap^14^.

We further propose that the ground-state should only be removed from the ensemble model if the occupancy of the refined ground-state conformer is ⪅10% – only in this case is the benefit of the ground-state model in refinement likely negligible. *We should assume that the ground-state is present in the ligand-bound crystal until it is proven absent*; this is contrary to the current convention, which appears to assume the opposite.

Correct parameterisation of the ensemble model can lead to complicated models and refinement constraints that are currently not supported by some refinement programs (REFMAC^15^, phenix.refine^11^): in some cases not shown here, we have found that refinement of multiple conformer models permitted occupancies for amino acids that summed to greater than unity. Further work will be required to generate occupancy and structural restraints that allow complex ensemble refinement in the general modelling case, without permitting unphysical atomic models. Procedural generation of ensembles and the corresponding parameterisation files will be critical to the uptake of this approach.

The examples shown here also highlight that RSCC alone is not enough to assess the quality of a ligand model: RSZD and RSZO should be used to ensure things have been modelled correctly, but require phases to be near convergence; a small B-factor ratio indicates consistency with the ligand’s environment; and a small RMSD measures stability in refinement. The combination of a normalised RSZO and B-factor ratio further allow the stability of B-factor and occupancy refinement to be analysed; imbalances between these two metrics are a good indication of imbalance in the occupancy and the B-factors. The radar plots present the validation metrics clearly, and may be a useful tool for the validation of ligands in general. In this manuscript, we have used the validation plots to compare multiple models, and to this end, the plot axes were re-scaled to cover the range of the data. However, we propose that a more general use of the radar plot is to show when the ligand scores depart from ideal values (the proposed ranges for the metrics are shown in section S2); examples are shown in Figure 6 for the ligand in section 4.3.

**Figure 6.**
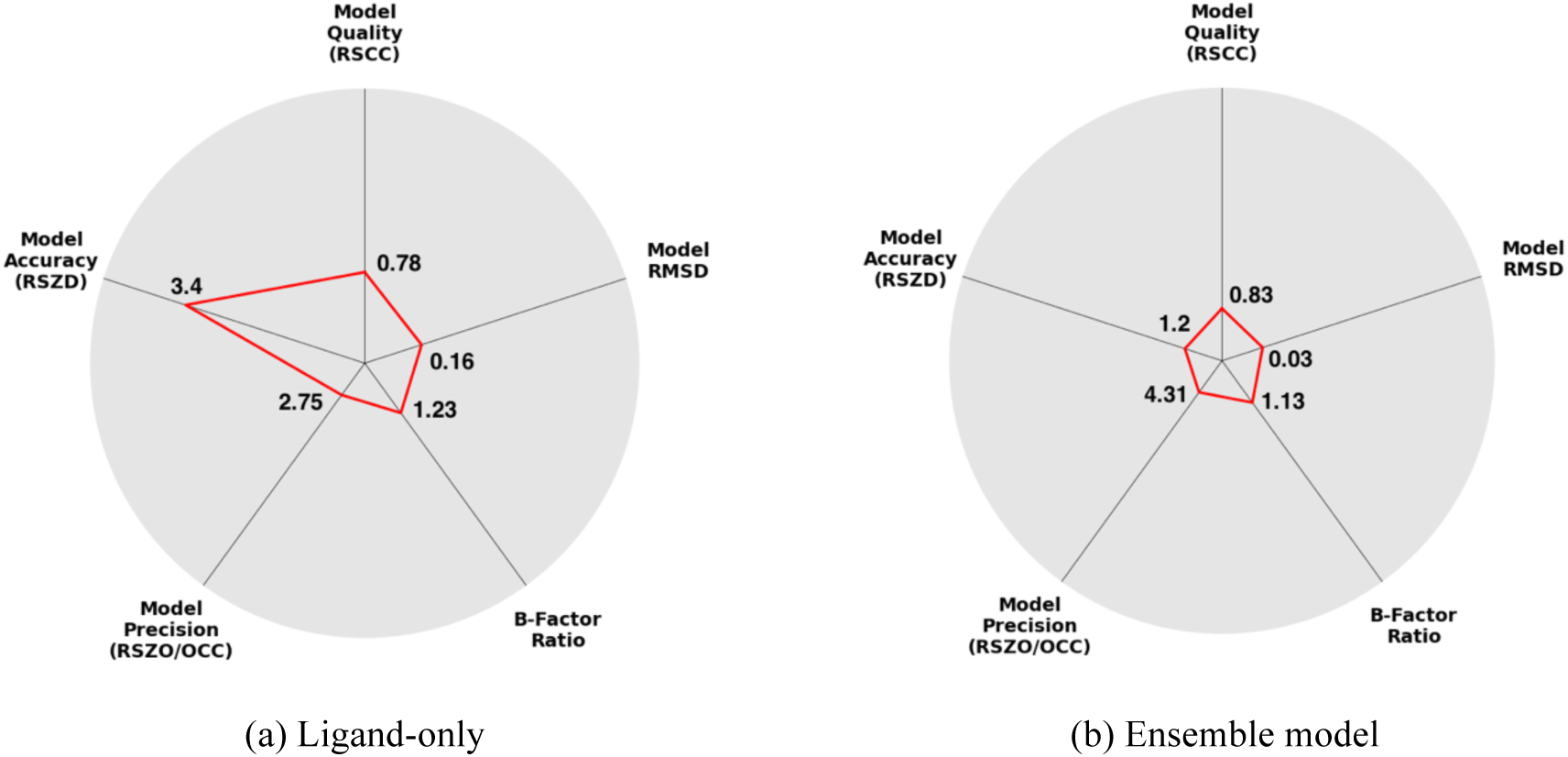
Radar plots clearly display the suspect features of a ligand, and indicate when validation scores deviate from ideal values. Validation plots for the ligand in section 4.3: (a) for the ligand-model only and (b) for the ligand when refined as an ensemble (scores are for the ligand residue only). Limits and thresholds for the validation plots are detailed in Table S5. The ligand-only model shows that un-modelled features are present, with a large RSZD. The ensemble model (with high-quality phases) scores well on all metrics, and remains close to the centre of the plot.

Lastly, we have investigated the impact of phase degradation on ligand model quality, compared to the effect of local modelling. We conclude that the modelling of local ground-state atoms is generally far more important than convergence of the global model, especially as global errors in typical modelling situations are likely to be much less than the ~30° phase error introduced here. “Tweaking” of sidechain conformations and water molecules in distant regions in the model to improve phases is likely not of importance if the binding of a ligand is the feature of interest. However, the modelling of the environment around and “under” the ligand is conversely of great importance.

Recent reports have emphasised the importance of achieving maximally correct phases for more reliable interpretation of weak difference density^13^. Instead, this work indicates that the main rationale for doing so is to ensure the validation metrics are reliable – as ligand identification can be performed without optimal phases^5^ – and that the refinement of occupancy and B-factors is stable (as demonstrated in section 4.4).

## Acknowledgements

NMP would like to recognize funding from EPSRC grant EP/G037280/1, UCB Pharma and Diamond Light Source. Crystallographic data were collected by Anthony Bradley, Patrick Collins, Romain Talon and Tobias Krojer at Diamond Light Source, Beamline i04-1. The SGC is a registered charity (No. 1097737) that receives funds from AbbVie, Bayer, Boehringer Ingelheim, the Canada Foundation for Innovation, the Canadian Institutes for Health Research, Genome Canada, GlaxoSmithKline, Janssen, Lilly Canada, the Novartis Research Foundation, the Ontario Ministry of Economic Development and Innovation, Pfizer, Takeda and the Wellcome Trust (092809/Z/10/Z).

## Supporting information

### S1. Crystallographic Information for Examples

The crystallographic parameters for each of the models used in the examples are listed in Table S1–Table S4. All models are refined with phenix.refine^11^ using the standard settings. All phase differences are calculated with cphasematch^10^ from the model phases as output by phenix.refine, relative to the ensemble-model phases. Occupancy refinement was performed on all models, except for the ground-state-only model. When multiple conformations were modelled, the occupancies are constrained to sum to unity.

**Table S1.** Crystallographic parameters and ligand model scores for the model in Section 4.2.

| Model | Mean Phase<br>Diff. (°) | R-work/<br>R-free | Occ | RSCC | RSZD | RSZO/<br>OCC | B-factor<br>Ratio | RMSD<br>(Å) |
| --- | --- | --- | --- | --- | --- | --- | --- | --- |
| Solvent Only | 2.92 | 0.129 / 0.171 | - | n/a | n/a | n/a | n/a | n/a |
| Ligand Only | 9.97 | 0.147 / 0.195 | 0.79 | 0.44 | 4.3 | 0.76 | 3.54 | 0.26 |
| Ensemble | - | 0.127 / 0.171 | 0.26 | 0.87 | 0.5 | 5.38 | 1.47 | 0.01 |
| Degraded Ensemble | 24.17 | 0.241 / 0.290 | 0.27 | 0.77 | 1.0 | 4.81 | 1.40 | 0.05 |

**Table S2.** Crystallographic parameters and ligand model scores for the model in Section 4.2.

| Model | Mean Phase<br>Diff. (°) | R-work/<br>R-free | Occ | RSCC | RSZD | RSZO/<br>OCC | B-factor<br>Ratio | RMSD<br>(Å) |
| --- | --- | --- | --- | --- | --- | --- | --- | --- |
| Solvent Only | 2.95 | 0.183 / 0.217 | - | n/a | n/a | n/a | n/a | n/a |
| Ligand Only | 4.15 | 0.184 / 0.215 | 0.68 | 0.92 | 3.6 | 1.62 | 1.41 | 0.20 |
| Ensemble | - | 0.182 / 0.216 | 0.41 | 0.96 | 1.6 | 3.41 | 1.04 | 0.02 |
| Degraded Ensemble | 31.06 | 0.311 / 0.363 | 0.29 | 0.90 | 0.1 | 1.72 | 1.18 | 0.70 |

**Table S3.** Crystallographic parameters and ligand model scores for the model in Section 4.3.

| Model | Mean Phase<br>Diff. (°) | R-work/<br>R-free | Occ | RSCC | RSZD | RSZO/<br>OCC | B-factor<br>Ratio | RMSD<br>(Å) |
| --- | --- | --- | --- | --- | --- | --- | --- | --- |
| Solvent Only | 3.86 | 0.157 / 0.219 | - | n/a | n/a | n/a | n/a | n/a |
| Ligand Only | 4.38 | 0.164 / 0.222 | 0.80 | 0.78 | 3.40 | 2.75 | 1.23 | 0.16 |
| Ensemble | - | 0.159 / 0.220 | 0.51 | 0.83 | 1.20 | 4.31 | 1.13 | 0.03 |
| Degraded Ensemble | 28.48 | 0.274 / 0.332 | 0.59 | 0.71 | 0.50 | 2.54 | 1.19 | 0.17 |

**Table S4.** Crystallographic parameters and ligand model scores for the model in Section 4.4.

| Model | Mean Phase<br>Diff. (°) | R-work/<br>R-free | Occ | RSCC | RSZD | RSZO/<br>OCC | B-factor<br>Ratio | RMSD<br>(Å) |
| --- | --- | --- | --- | --- | --- | --- | --- | --- |
| Solvent Only | 4.22 | 0.186 / 0.216 | - | n/a | n/a | n/a | n/a | n/a |
| Ligand Only | 2.20 | 0.183 / 0.213 | 0.89 | 0.95 | 2.00 | 4.38 | 1.26 | 0.03 |
| Ensemble | - | 0.182 / 0.212 | 0.84 | 0.96 | 1.60 | 5.95 | 1.20 | 0.01 |
| Degraded Ensemble | 34.46 | 0.341 / 0.380 | 0.77 | 0.91 | 0.10 | 3.77 | 1.14 | 0.07 |

### S2. Validation Radar Plots

Standard validation plots are generated by recording the density scores radially on the graph axes and connecting these points with lines. For the comparative plots (Figure 4), the axes are re-scaled such that the limits are the minimum and maximum of the metric scores. For normal validation plots (Figure 6), the limits of each of the scores are shown in Table S5. These plots can be generated using the giant.score_model script distributed as part of the giant package within the panddas package, available as part of CCP4^10^.

**Table S5.** Radar plot axes limits. The limits and length scales for the radial axes are defined here. The inner limit defines the value at which the plotted line will begin to move away from the centre of the plot. The outer limit defines the values at which the plotted line will reach the end of the radial axis, and be plotted outside the graph area. If a metric is inverted, large values will be plotted closer to the centre of the radar plot, and smaller values will be plotted further from the centre.

| Metric | Inner Limit | Outer Limit | Inverted |
| --- | --- | --- | --- |
| RSCC | 0.85 | 0.6 | Yes |
| RSZD | 1.5 | 4 | No |
| RSZO/OCC | 2 | 0 | Yes |
| B-factor Ratio | 1 | 3 | No |
| RMSD | 0 | 1.5 | No |

**Figure S1.**
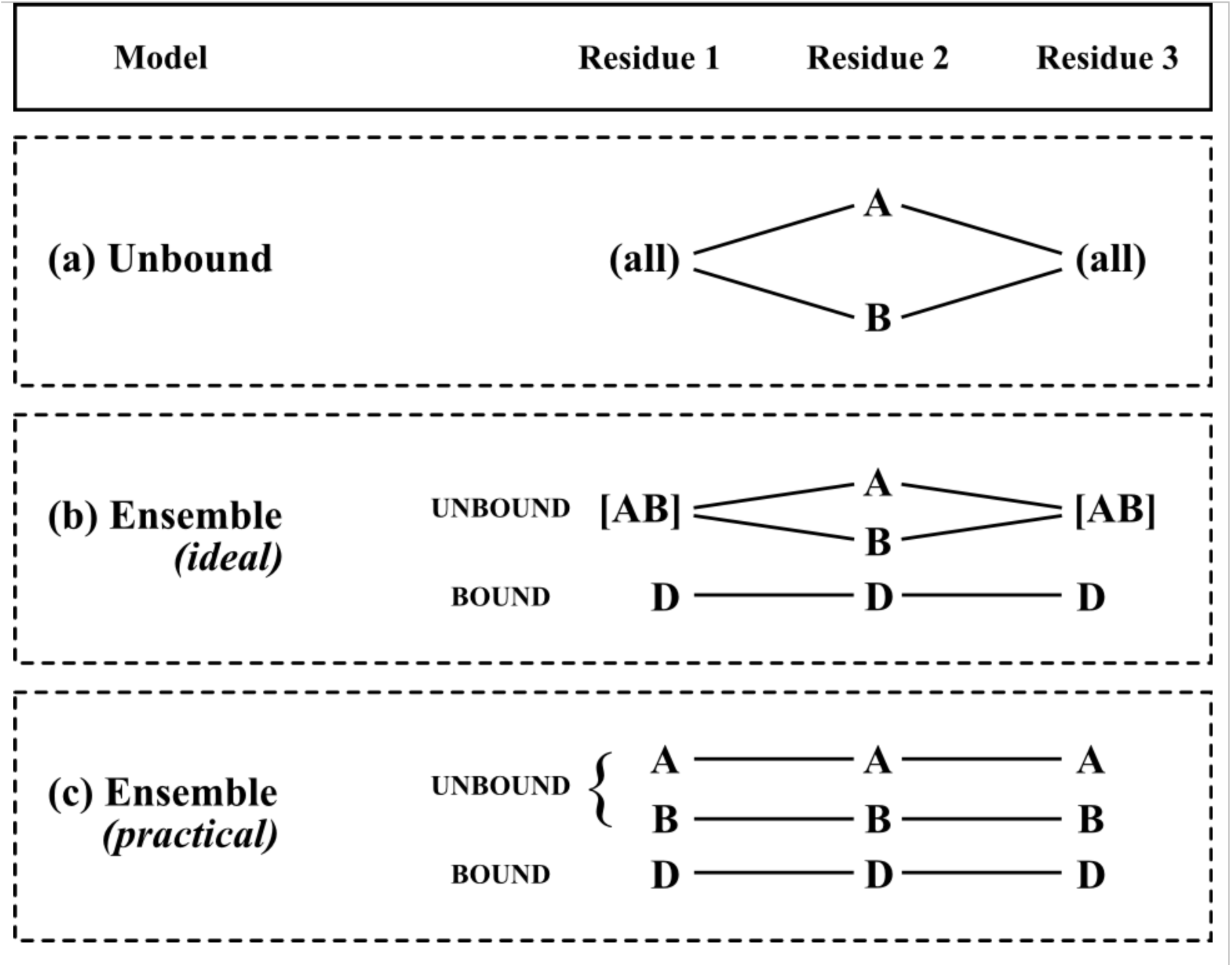
Limitations of alternate conformers require workarounds in modelling. Conformers in crystallographic models do not support branching of conformers – where alternate sub-conformations can be added to existing alternate conformations – so workarounds are required where complex alternate conformations are present. (a) The unbound structure contains a residue in alternate conformations. These are assigned to conformers A and B. (b) In the ideal case, a new conformer (D) could be introduced without editing the existing structure. However, it is not possible to create the [AB] conformer (conformers must be either A *or* B). If the [AB] conformation is labelled as A, then the B-conformation of residue 2 becomes disconnected in refinement. (c) The workaround is to enumerate all conformations of [AB] for each of the surrounding residues; this creates multiple conformations A and B of residue 1 and residue 3.

